# In-cell cryo-electron tomography reveals differential effects of type I and type II kinase inhibitors on LRRK2 filament formation and microtubule association

**DOI:** 10.64898/2025.12.18.694444

**Authors:** Tamar Basiashvili, Joshua Hutchings, Siyu Chen, Eva P. Karasmanis, W. Alexander Flaherty, Andres E. Leschziner, Elizabeth Villa

## Abstract

Mutations in Leucine-Rich Repeat Kinase 2 (LRRK2) are a leading contributor to developing familial and idiopathic Parkinson’s disease (PD). Most PD-causing LRRK2 mutations increase the kinase activity, leading to increased phosphorylation of Rab GTPases, disrupting vesicular trafficking, cytoskeletal dynamics, and autophagy. Under homeostatic conditions, the bulk of WT and PD-mutant LRRK2 is found in the cytosol. However, exogenously expressed LRRK2 can form microtubule-associated filaments that have been shown to affect molecular transport along microtubules in vitro. While the physiological relevance of microtubule binding has not been established yet, inhibitors being designed and tested as therapeutics have been shown to either promote or prevent filament formation of LRRK2. In this study, we examine the localization and resulting molecular organization of hyperactive LRRK2-I2020T, a common PD mutant, in cells treated with type I (MLi-2) or type II (GZD-824) kinase inhibitors. Treatment with a type I kinase inhibitor results in extensive LRRK2-I2020T decoration around microtubules and microtubule bundling. Stabilization of LRRK2-I2020T filaments by type I inhibitor treatment allowed us to build a full-length closed-kinase model of LRRK2-I2020T in its cellular environment. Conversely, treatment with a type II inhibitor resulted in minimal microtubule decoration by LRRK2-I2020T compared to Type I inhibitor treated cells. This study provides a structural framework for understanding how type I and type II kinase inhibitors differentially modulate LRRK2 filament formation, demonstrating that type I inhibitor treatment promotes a distinct filament architecture, whereas such assemblies are not observed with type II inhibitors.

## Introduction

Parkinson’s disease (PD) is a chronic and progressive neurodegenerative disorder affecting approximately ten million people worldwide^1–4^. Genome-wide association studies have identified leucine-rich repeat kinase 2 (LRRK2) as a major risk factor for developing familial and idiopathic PD^5–7^. LRRK2 contributes to disrupted cytoskeletal dynamics, lysosomal dysfunction, neuroinflammation and impaired dopaminergic signaling^8–11^.

LRRK2 is a large multidomain protein, one in four in the human genome to contain a GTPase and a kinase^12–14^. Its C-terminal catalytic region is composed of ROC GTPase, COR, Kinase and WD40 (RCKW) domains (Fig. 1A). Pathogenic mutations in this region (e.g. I2020T, G2019S, R1441H/G/C/S, Y1699C) increase the kinase activity leading to hyperphosphorylation of Rab GTPases disrupting fusion, docking and trafficking of vesicular membranes^15–18^. The N-terminal region mediates protein-protein interactions, facilitating recognition of LRRK2 substrates through the Leucine rich repeats (LRR), Ankyrin (ANK) and Armadillo (ARM) domains (Fig. 1A). In the autoinhibited inactive kinase state, the N-terminus occludes catalytic domains and limits substrate access to the kinase domain^19–21^. Upon activation, these N-terminal repeats are proposed to undock, enabling substrate binding near the kinase moiety and subsequent phosphorylation of the target^19^. Recent structural studies have elucidated the architecture of LRRK2’s catalytic RCKW domains in both active and inactive conformations, as well as full-length autoinhibited and intermediate activation state structures of the protein^19–27^. Resolving the full-length closed-kinase structure of LRRK2 remains a significant challenge due to the large conformational heterogeneity of the N-terminal repeats in the active state.

**Figure 1.**
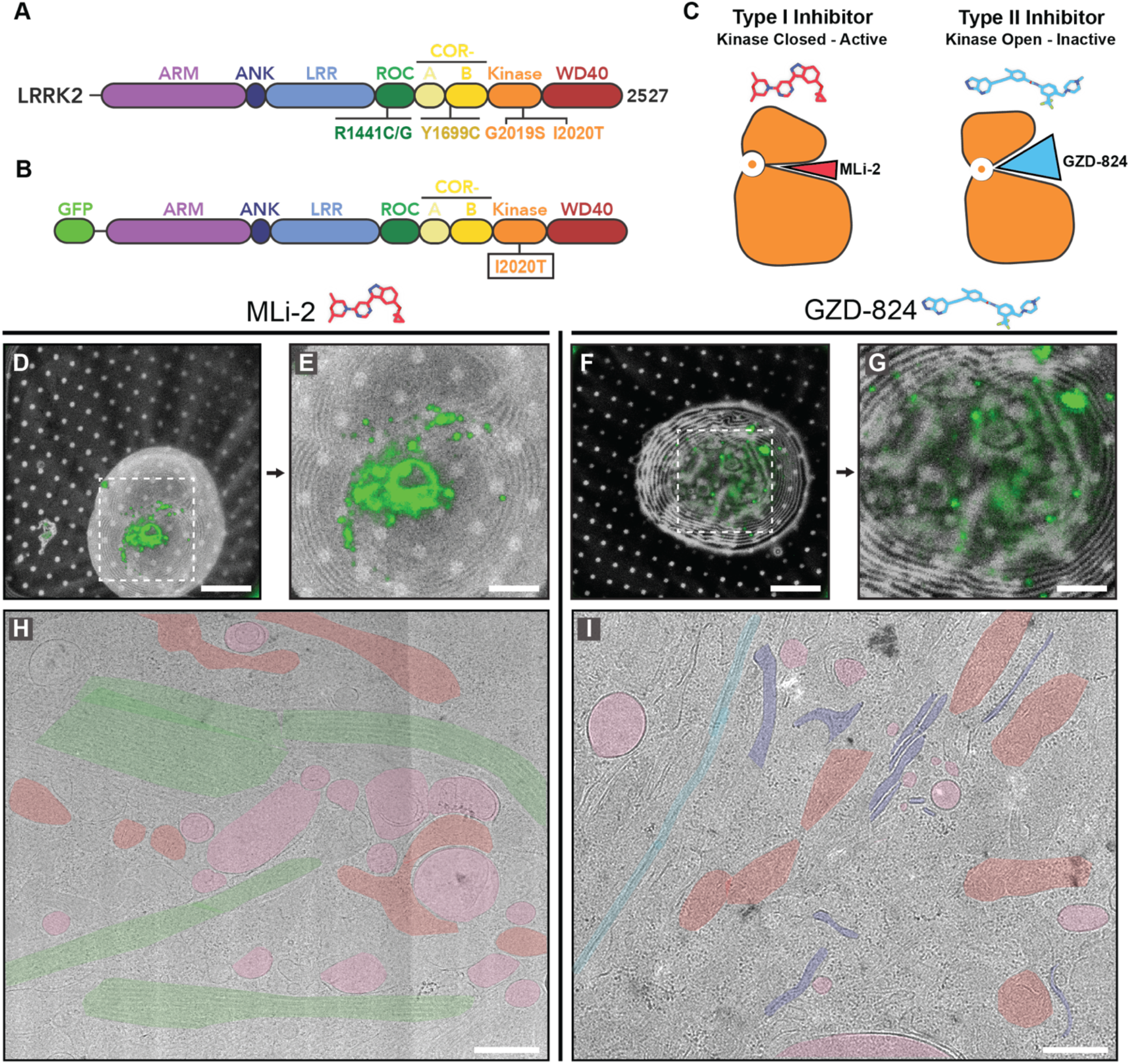
LRRK2^IT^ expression in 293T cells responds differently to treatment with type I and type II kinase inhibitors. **(A)** LRRK2 contains multiple functional domains including Armadillo (ARM), Ankyrin repeats (ANK), Leucine-rich repeats (LRR), Ras of complex (ROC), COR, kinase and WD40 domain. **(B)** Schematic of the LRRK2 protein construct used in this paper that includes a hyperactive I2020T mutation and an N-terminal GFP-tag. **(C)** Type I inhibitors (red) capture the LRRK2 kinase in active, closed conformation while type II inhibitors (blue) trap kinase of LRRK2 in an inactive, open kinase conformation. **(D-E)** Cryo-fluorescence microscopy (cryo-FM) image of a vitreous cell on an electron microscopy grid expressing LRRK2^IT^ treated with MLi-2. The GFP-LRRK2^IT^ signal (green) is present as filaments as well as puncta. **(F-G)** Cryo-FM image of a cell expressing LRRK2^IT^ treated with GZD-824. The GFP-LRRK2^IT^ signal (green) shows a punctate distribution of the protein throughout the cytosol of the cell. **(H-I)** Cryo-TEM overview of the lamella of a cell expressing LRRK2^IT^ treated with MLi-2 (H) and GZD-824 (I). Microtubule bundles are highlighted in green with some displaying dense LRRK2 decoration. Green: microtubule bundles, red: mitochondria, pink: vesicles, blue: plasma membrane, purple: endoplasmic reticulum; Scale bars: D, F 20μm; E, G 5μm; H, I 500nm.

Given its role in PD pathogenesis, LRRK2 has been the focus of small molecule kinase inhibitor development with two drug candidates currently in clinical testing^28–31^. There are two major classes of LRRK2 inhibitors: type I LRRK2 specific inhibitors (e.g. MLi2, LRRK2-IN-1, HG-10-102-01) stabilize the kinase domain in its closed active-like conformation, while type II broad spectrum inhibitors (e.g. GZD-824, ponatinib, rebastinib) stabilize the kinase in its open, inactive-like conformation^31–37^ (Fig. 1C). Both types of inhibitors bind to the kinase domain, but type I inhibitors compete for ATP binding while type II inhibitors bind to the allosteric site adjacent to the kinase domain^19,21^. The conformational changes of the kinase domain when treated with different types of inhibitors have been characterized, but the effect of inhibitor treatment on LRRK2 localization and molecular architecture in cells remains unclear^19,21^.

In cells, LRRK2 is typically cytosolic, but under certain conditions it binds to microtubules and forms extensive filaments^22,38,39^. Association with microtubules disrupts molecular transport along microtubules in vitro^23^.

Hyperactivating pathogenic mutations such as I2020T and R1441H/G/C/S increase the protein’s propensity to form microtubule associated filaments^22–24^. Additionally, type I, but not type II, kinase inhibitor treatment increases LRRK2 filamentation in cells^24,40^. While physiological significance of LRRK2 filamentation remains unclear, microtubule association provides an assay to test effects of kinase inhibitors on molecular architecture of LRRK2 in cellular models.

In this study, we investigate the effects of different types of kinase inhibitors on the cellular architecture and localization of PD-associated LRRK2-I2020T (LRRK2^IT^). To this end, we expressed GFP-tagged LRRK2^IT^ in 293T cells and treated samples with either type I (MLi-2) or type II (GZD-824) inhibitors. To visualize effects of inhibitor treatment on LRRK2 architecture at nanometer resolution we used cryo-correlative light and electron microscopy (cryo-CLEM), which combines cryo-electron tomography with cryo-fluorescence microscopy (cryo-FM)^41–43^.

In cells treated with MLi-2, we observed highly ordered LRRK2-decorated microtubules that formed extensive microtubule bundles of 15-30 microtubules. In contrast, LRRK2 microtubule association was rarely observed in GZD-824 treated cells. These observations point to distinct mechanisms by which type I and type II kinase inhibitors influence LRRK2 filamentation in a cellular context. Besides increased filament formation, MLi-2 treatment reduced conformational heterogeneity of LRRK2^IT^, allowing us to characterize the architecture of the full-length microtubule-associated conformation of LRRK2^IT^. We present the structure of full-length LRRK2^IT^ in a closed-kinase conformation and N-terminal repeats undocked from the RCKW domains, providing insight about hyperactive protein architecture in the cellular environment.

## Results

### LRRK2^IT^ localizes to microtubules in the presence of a type I inhibitor and to the cytosol with a type II inhibitor

We used cryo-CLEM to study how cellular localization of GFP-tagged hyperactive LRRK2^IT^ is affected by treatment with either type I (MLi-2) or type II (GZD-824) inhibitors (Fig. 1; Ext. Fig. 1). Our previous work showed that expression of LRRK2^IT^ in 293T cells forms an extensive network of filaments around microtubules^22^. Here we added either MLi-2 or GZD-824 inhibitors that resulted in significant phenotypic differences in LRRK2^IT^ localization (Fig. 1D-G; Ext. Fig. 1A-J). In cells treated with MLi-2, we observed LRRK2^IT^ in extended filaments, puncta, and diffuse in the cytosol (Fig. 1D-E; Ext. Fig 1A-D). In contrast, when cells were treated with GZD-824, LRRK2^IT^ was mostly localized to puncta and distributed throughout the cytosol, with reduced filament formation (Fig. 1F-G; Ext. Fig 1E-H), in agreement with our previous work^23,24,40^.

To probe this difference at the structural level, we used cryo-FIB milling to produce thin samples of vitrified cells for cryo-ET. Cryo-FM images were aligned to cryo-TEM overviews of the lamella to localize LRRK2^IT^ (Fig. 1H-I; Ext. Fig. 1I-J). In cells treated with MLi-2, we co-localized LRRK2^IT^ with an extended network of microtubules (Fig. 1H-I; Ext. Fig. I-J). These microtubules were coated with a dense LRRK2^IT^ lattice and bundled. The LRRK2^IT^ decorated microtubule network was extensive, spanning 2.17±1.40µm in length (Fig. 1H; Ext. Fig. 1K). By contrast, microtubule bundling was not present in GZD-824 treated cells (Fig. 1I).

### Type I kinase inhibitor treatment promotes bundling of LRRK2^IT^ decorated microtubules

We acquired cryo-electron tomograms of MLi-2 and GZD-824 treated cells targeting LRRK2^IT^ localization sites and identified all microtubules within the data. In MLi-2 treated cells, we focused on microtubules that were decorated with a LRRK2^IT^ lattice (Fig. 2A-D; Ext. Fig. 2A-B; Ext. Fig 3). We found a substantial fraction of them to be surrounded by other decorated microtubules often closely spaced and arranged in parallel, suggesting that LRRK^IT^ binding promotes microtubule bundling (Fig. 2A-D; Ext. Fig. 2A-B). Bundles of LRRK2^IT^ decorated microtubules contained an average of 16±8.04 microtubules (range 5-30; Fig. 3M). The center-to-center distance between LRRK2^IT^-decorated microtubules was 68.7nm±5.73nm, while undecorated microtubule bundles had a center-to-center distance of 35nm±3.00nm in our data and in previous findings (Fig. 3L)^22^. The expanded spacing observed in the LRRK2^IT^ decorated microtubules is due to the 15nm thick LRRK2^IT^ layer that coats each microtubule, adding 30nm to the center-to-center distance (Fig. 3A-C, L).

**Figure 2.**
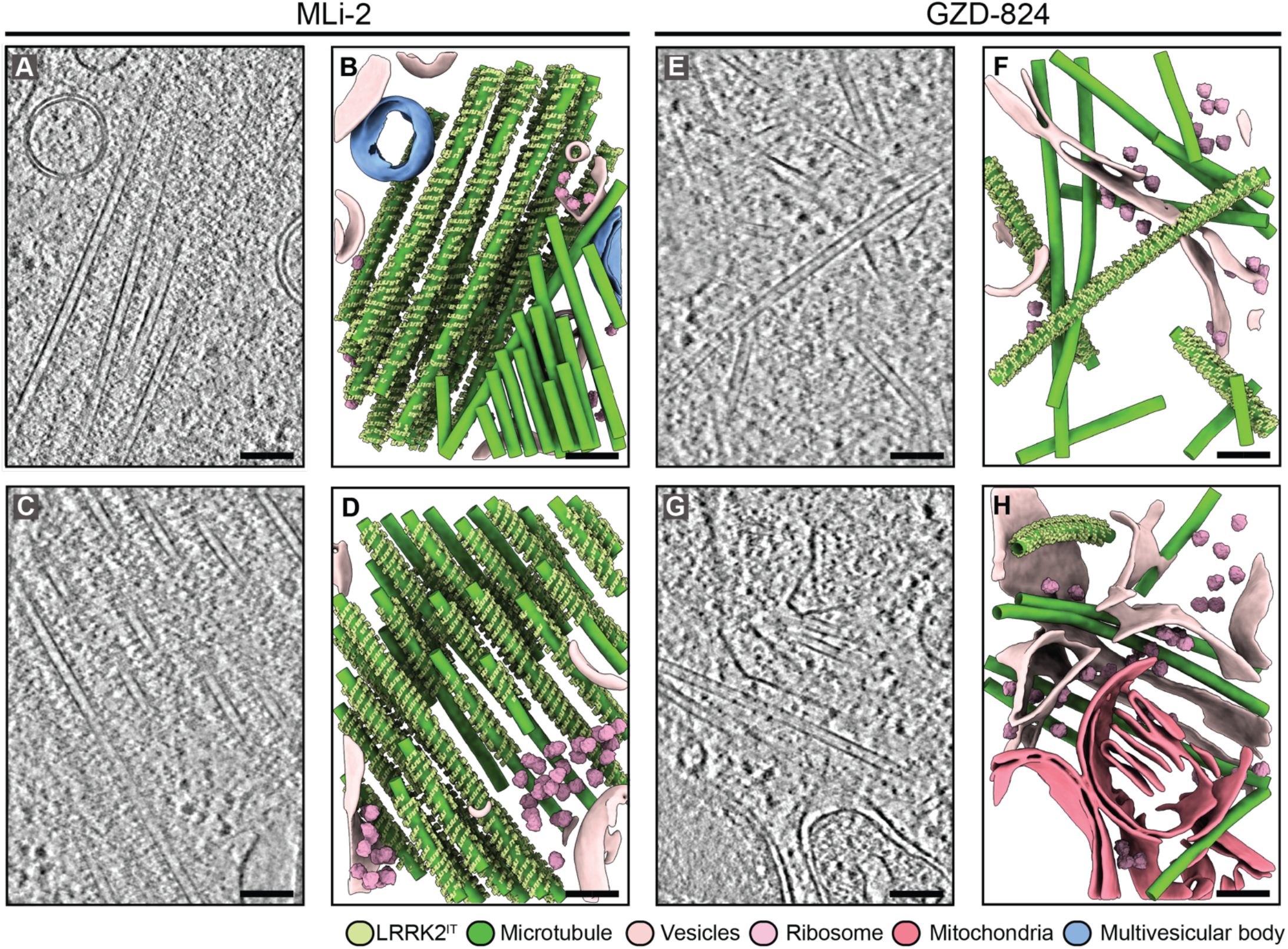
Treatment with the type I kinase inhibitor MLi-2 promotes formation of higher-order LRRK2^IT^-decorated microtubule bundles, whereas the type II kinase inhibitor GZD-824 reduces LRRK2^IT^ filamentation and microtubule bundling in cells. **(A, C)** Representative tomogram slices through the cytosol of a cell treated with MLi-2, showing a bundle of microtubules decorated with LRRK2^IT^. **(B, D)** Segmentation of the tomogram in A, C highlighting bundles of microtubules decorated with a well-ordered LRRK2^IT^ lattice. **(E)** Tomogram slice of a cell treated with GZD-824, showing sparse LRRK2^IT^ decoration on microtubules. **(F)** Segmentation of the tomogram in E. LRRK2^IT^ decoration is present on three microtubules while there are nine undecorated microtubules present in the proximity. **(G)** Representative image of a cell treated with GZD-824. Single microtubule is decorated with LRRK2^IT^. This distribution pattern contrasts sharply with the extensive microtubule-associated filaments observed following Type I inhibitor treatment. **(H)** Segmentation of a tomogram in G. of a cell treated with GZD-824. Six undecorated microtubules are present near a single LRRK2 decorated microtubule. Lime Green: LRRK2^IT^, green: microtubule, red: mitochondria, light pink: vesicles, blue: multivesicular bodies, pink: ribosomes. Scale bars: 100nm.

**Figure 3.**
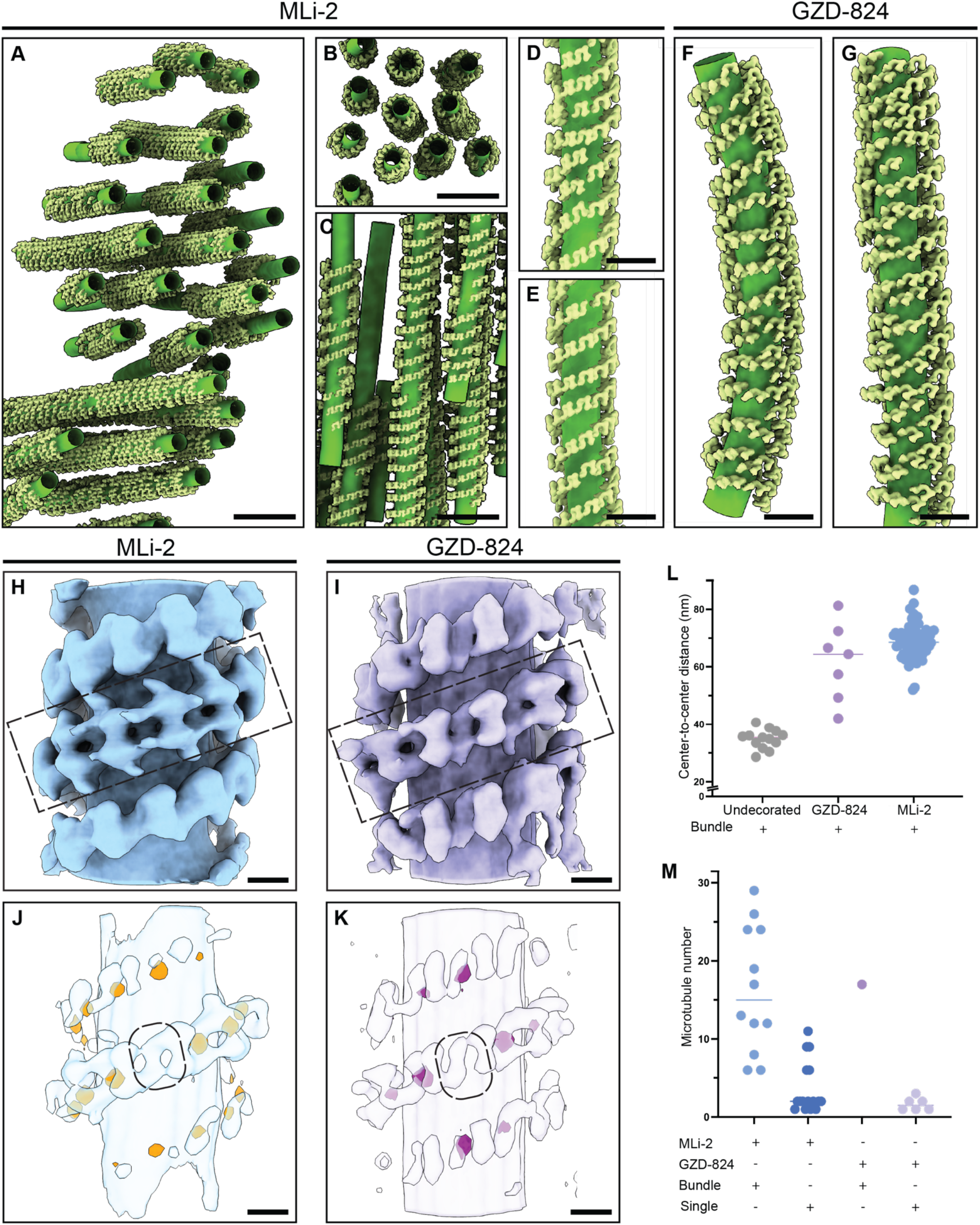
LRRK2^IT^ lattice that decorates microtubules is highly ordered in cells treated with type I as compared to type II inhibitor. **(A)** Segmented representation of a LRRK2^IT^ associated microtubule bundle in MLi-2 treated cells. Individual LRRK2^IT^ subunits are highly ordered around the microtubules. **(B)** Top-down view of the segmented representation of a LRRK2^IT^ associated microtubule bundle in MLi-2 treated cells. The individual microtubules are separated from each other by 65-70nm. **(C)** Segmented representation of a LRRK2^IT^ associated microtubule bundle in MLi-2 treated cells. **(D-E)** Close-up view of the LRRK2^IT^ decoration on microtubules in MLi-2 treated cells. **(F-G)** Segmented representation of LRRK2^IT^ associated microtubules in GZD-824 treated cells. LRRK2^IT^ lattice is less ordered compared to MLi-2 treated lattice in (D-E). *(continued)* **(H-I)** Subtomogram average of the LRRK2^IT^ associated with microtubules from MLi-2 (H) or GZD-824 (I) treated cells. MLi-2 treated cells exhibit a more ordered lattice of LRRK2^IT^ than the lattice found in GZD-824 treated cells. **(J-K)** Nearest neighbor analysis of a central LRRK2^IT^ subunit (highlighted in black) in MLi-2 and GZD-824 treated cells. In MLi-2 treated cells, the central LRRK2^IT^ subunit has 16 nearest neighbors highlighted in orange. While in GZD-824 treated cells, the central LRRK2^IT^ subunit has only 8 nearest neighbors shown in pink. **(L)** Evaluation of the center-to-center distance of undecorated and LRRK2^IT^-decorated microtubule bundles in MLi-2 and GZD-824 treated cells. (MLi-2 n=80, GZD-824 n=7, undecorated n=13; vertical line at median). **(M)** Quantification of LRRK2^IT^ decorated microtubule (MT) bundles and isolated microtubules in MLi-2 and GZD-824 treated cells. (bundles quantified in MLi-2 treated cells n=12, isolated MTs quantified in MLi-2 treated cells n=16, isolated MTs quantified in GZD-824 n=6, bundle of MTs quantified in GZD-824 bundle n=1, vertical line at median). Scale bars: A-C 40nm; D-G 20nm; H-K 10nm.

In contrast, tomograms acquired from type II GZD-824 inhibitor treatment resulted in reduced LRRK2 filamentation around microtubules, with only a few rare, isolated microtubules showing LRRK2^IT^ decoration (Fig. 2E-H; Ext. Fig. 2C-D). Interestingly, we captured a single bundle of LRRK2^IT^ decorated microtubules in the tomograms of GZD-824 treated cells (Ext. Fig. 4). This bundle contained 15 microtubules with intermicrotubule spacing similar to LRRK2^IT^ decorated microtubule bundles in MLi-2 treated cells. Otherwise, fluorescence microscopy of GZD-824 treated cells showed LRRK2^IT^ was present as discrete cytosolic puncta, suggesting association with intracellular membranes (Fig. 1F-G, I; Ext. Fig. 1E-H, J). In these data, organelles appeared intact and morphologically unaltered, with no discernible densities, or organized assemblies that could be confidently attributed to LRRK2^IT^ (Fig. 1I). Aside from the absence of microtubule bundling, the overall ultrastructure of GZD-824–treated cells closely resembled that of MLi-2–treated and of untreated cells^22^.

**Figure 4.**
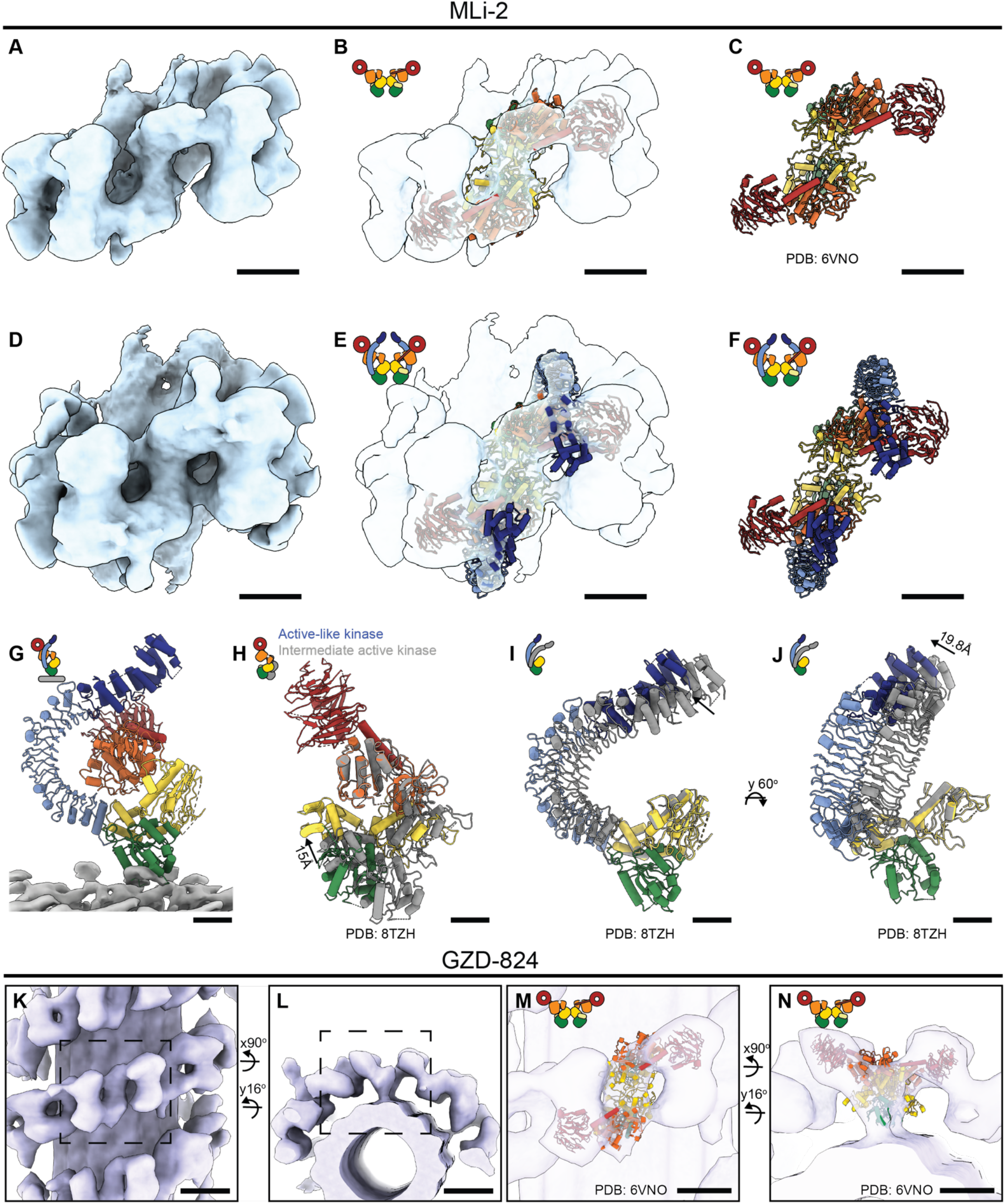
Subtomogram analysis of LRRK2 MLi-2 reveals structural details about a N-terminal domains of closed-kinase LRRK2^IT^. (A) Subtomogram average map of the microtubule-bound LRRK2 from MLi-2 treated cells (threshold value 0.24). **(B-C)** Molecular model of two central protomers of active kinase LRRK2 RCKW fit into map A. The protomers of LRRK2 fit well within the density corresponding to the active conformation of the LRRK2 kinase. The domains are colored according to Fig. 1A. **(D)** Subtomogram average of LRRK2 displaying densities for the N-terminal domains emanating from the catalytic half of the protein (threshold value 0.14). **(E-F)** Molecular model of the two central protomers of full-length LRRK2 fit into the map shown in D. The model includes RCKW domains and N-terminal LRR-ANK domains. **(G)** Molecular model of a single subunit of full length LRRK2 in active conformation interacting with a microtubule (gray) via ROC domain (green). *(continued)* **(H)** Comparison between the RCKW domains of LRRK2 in intermediate active kinase conformation (gray) and fully active full-length in-cell model of LRRK2 (colored). The kinase domain in the cellular map is fully closed, demonstrated by 15Å shift between the COR domains. **(I-J)** Comparison of the LRR and ARM domains of LRRK2 in the in-cell model (colored) versus the intermediate kinase-active LRRK2 structure (gray). The COR– ROC domains are aligned in both structures to highlight the positional shifts in the LRR and ARM domains **(K-L)** Subtomogram average of microtubule-bound LRRK2 from GZD-824–treated cells. **(M-N)** Molecular model of the active kinase RCKW domains fitted into the map shown in panel K. The molecular model aligns well with the active kinase conformation of LRRK2. The N-terminal domains are not visible in this map. Rotated views are shown. Scale bars: A-F 5nm; G-J 2nm; K-L 10nm; M-N 5nm.

### LRRK2^IT^ microtubule decoration is highly ordered in cells treated with type I compared to type II inhibitors

To characterize the differences between LRRK2^IT^ lattice organization in cells treated with different inhibitors, we performed subtomogram analysis of decorated microtubules and mapped the location of each individual LRRK2 dimer and quantified lattice parameters (Fig. 3). In the MLi-2 treated cells, LRRK2^IT^ strands were organized around microtubules with a regularly spaced lattice, similar to the LRRK2^IT^ strands in cells not treated with the inhibitor (Fig. 3A-E)^22^. Conversely, in GZD-824-treated cells, the LRRK2^IT^ lattice was rare, and when existent, it was disordered (Fig. 3F-G). While average pitch, rise, and handedness of the filaments of the rare GZD-824 treated LRRK2 filaments were similar between the MLi-2 parameters (Fig. 3H-K; Ext. Fig. 5), the relative orientation of individual dimers was variable, and the lattice was highly irregular of in GZD-824 treated cell (Fig. 3F-G). Using nearest neighbor analysis, we found that in the LRRK2^IT^ lattice from MLi-2-treated cells, up to 16 close neighbors were placed with high precision, compared to only eight in the GZD-824-treated LRRK2^IT^ lattice (Fig. 3J-K; Ext. Fig. 5A-D). Altogether this suggests that MLi-2 treatment stabilizes LRRK2 filamentation on microtubules, while GZD-824 disrupts LRRK2^IT^ lattice weakening its association with microtubules.

**Figure 5.**
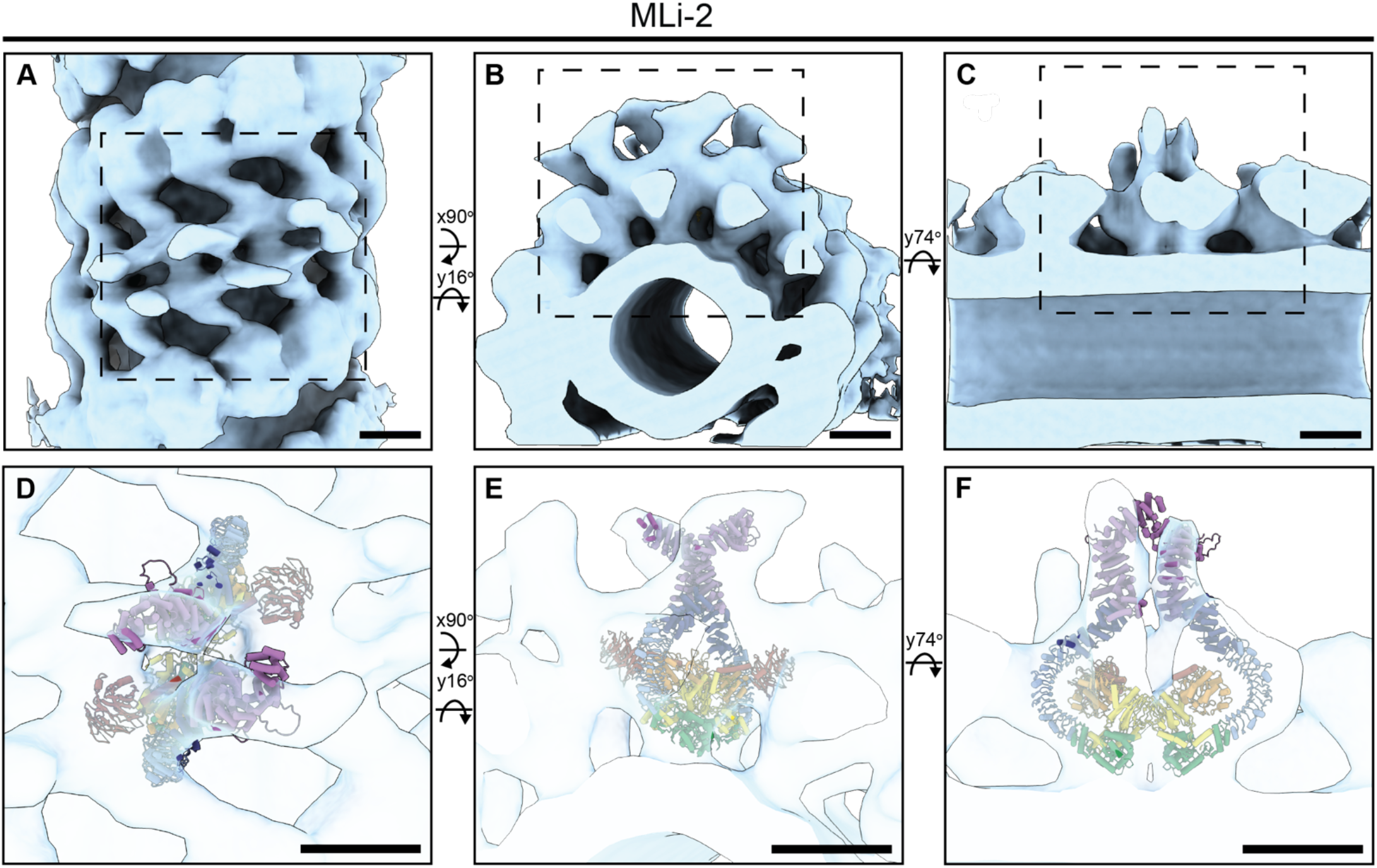
Model of full-length LRRK2^IT^ associated with microtubules in its active-like conformation. **(A)** Subtomogram average of LRRK2^IT^ lattice decorating a microtubule. **(B)** Cross-section view of the LRRK2^IT^ lattice. Catalytic domains are closest to the microtubule while outer layer represents the N-terminal protein-protein interaction domains of LRRK2. **(C)** Side view of the microtubule demonstrating the extension of the N-terminal layer of LRRK2. **(D-F)** Atomic model fit of full-length LRRK2^IT^ in the subtomogram average map. LRRK2 domain colors are as in Fig. 1A. Scale bars: A-C 10nm; D-F 10nm.

### Subtomogram analysis in type I inhibitor treated cells reveals the architecture of the N-terminal repeats of the full-length LRRK2 in closed-kinase conformation

Full-length LRRK2 structures with closed-kinase conformation have been determined, but the flexible N-terminal repeats remain disordered and are not resolved in these structures^19–27^. Given the stabilization of LRRK2^IT^ on microtubules with MLi-2 treatment we attempted to resolve full-length closed-kinase conformation of LRRK2^IT^ focusing on resolving the N-terminal repeat architecture.

First, we performed subtomogram analysis on regions of the microtubules that focused on a LRRK2^IT^ dimer and its surroundings (Fig. 4). We focused refinements on the COR-COR dimerization interface, centering the reconstruction around four LRRK2 monomers, resulting in a 12Å resolution map (map A; Fig. 4A). We also refined the WD40-WD40 dimerization interface and focused refinements on four central subunits reaching 16Å resolution (map B; Ext. Fig. 6). These reconstructions allowed us to dock all the individual domains of LRRK2 into the map (Fig. 4). To determine the conformation of the kinase domain of LRRK2^IT^ we fit the structures of active and inactive kinase of LRRK2 into map A (Fig. 4A-C). Fitting the inactive kinase structure resulted in significant clashes between COR-COR domains of neighboring LRRK2^IT^ subunits. The closed-kinase structure of the catalytic domain fit in map A (Fig. 4A) with no clashes, confirming that the reconstruction represents LRRK2^IT^ in its closed-kinase conformation (Fig. 4B-C). In maps A and B, we resolved a connection between a ROC domain and tubulin, present in the in-vitro microtubule-associated LRRK2 RCKW structure, but previously elusive in the in-situ maps (Fig. 4G, Ext. Fig. 6G-H)^23,24^.

Notably, our maps have ordered density for the N-terminal LRR-ANK domains (Fig. 4D-F), and thus we set out to build a full-length model of closed-kinase LRRK2^IT^. We based this model on our published structure of in-vitro MLi-2 treated full-length LRRK2, where the kinase was captured in an intermediate activation state with the N-terminal repeats docked to the kinase, preventing it from adopting a fully active conformation^19^. We split this model in three parts: the WD40 and C-lobe of the kinase, the N-lobe of the kinase with ROC and COR domains, and the LRR and ANK domains, aligned and fitted each of these three to our map A (Fig. 4D-F). Given the limited resolution of the map A, we fit the model as three rigid bodies without flexible fitting or atomic refinement. When comparing this structure with the reference model of LRRK2^IT^ in its intermediate active conformation, we discerned key differences between the structures. First, the kinase was in the fully active conformation in the in-situ map (Fig. 4G-H). Second, the LRR-ANK domains were undocked from the kinase and WD40 domains and shifted by 19.8Å as compared to the intermediate kinase-active structure (Fig. 4I-J). In the closed-kinase form, the N-terminal repeats are known to undock from the catalytic core (Fig. 4I-J), allowing easier access of LRRK2 substrates to the kinase domain. Here, we captured and modeled for the first time the N-terminal repeats undocked from the catalytic domains with kinase in an active-like conformation.

Additionally, we performed subtomogram analysis in Dynamo on a larger LRRK2^IT^-decorated lattice that contained three layers of LRRK2^IT^ density around the microtubule; we refer to this average as map C. Refinement was focused on the central four LRRK2^IT^ subunits to better resolve additional protein densities within this larger lattice. In map C (Fig. 5A; Ext. Fig. 7), we discerned an additional layer of protein decoration that accommodated not only LRR and ANK but also the ARM domain of LRRK2^IT^ (Fig. 5A-C; Ext. Fig. 7). In this map, we first fit in our in-situ model of LRRK2^IT^ and fit an AlphaFold2 model of the ARM domain into the additional densities (Fig. 5D-F; Ext. Fig. 7). While resolution of this map is moderate at 24.5 Å, densities that emanate from the catalytic part of the protein correspond to the LRR-ANK-ARM domains (Fig. 5D-F; Ext. Fig. 7; Ext. Fig. 8). LRR-ANK-ARM domains all point outward, extending away from the catalytic domains of LRRK2^IT^ (Fig. 5D-F). We hypothesize that the bundling visible in our maps could be reinforced by the connections between the LRRK2^IT^ N-terminal ARM domains between adjacent microtubules in a bundle (Fig. 3A-C; Fig. 5D-F).

Our attempts to characterize the molecular architecture of LRRK2^IT^ following treatment with GZD-824 showed minimal filamentation of LRRK2^IT^ on microtubules (Fig. 2E-H). In tomograms of GZD-824-treated cells, only six individual LRRK2-decorated microtubules and a single LRRK2 decorated microtubule bundle was detected (Fig. 2E-H, Ext. Fig. 4). Subtomogram analysis revealed a disrupted LRRK2 strand geometry, compared to that observed in MLi-2 treated cells (Fig. 3F-G). We obtained a subtomogram map of LRRK2^IT^ decoration at significantly lower resolution (∼30 Å), in which only the RCKW domains could be resolved (Fig. 4K–N). Fitting of the RCKW domains into this map was consistent with a closed-kinase conformation of LRRK2^IT^ (Fig. 4M–N). A likely explanation for this observation is that GZD-824 treatment was insufficient to convert all pre-existing, microtubule-bound LRRK2 molecules from an active to an inactive kinase conformation. Instead, the residual filaments detected are composed of LRRK2 molecules that formed prior to inhibitor addition and are refractory to subsequent GZD-824 binding. This interpretation is consistent with the markedly reduced frequency of filamentation observed under GZD-824 treatment.

## Discussion

Utilizing cellular structural biology tools, we dissected the effects of different types of inhibitors on the localization and architecture of hyperactive LRRK2^IT^ expressed in 293T cells. In cells treated with type I MLi-2 inhibitor, we captured a significant number of microtubules with LRRK2^IT^ decoration, the majority of which formed extensive bundled networks of LRRK2^IT^ filaments (Fig. 3). On the other hand, when cells were treated with GZD-824, only a few microtubules were decorated with LRRK2^IT^ (Fig. 2, E-H). Differences in LRRK2^IT^ propensity to form filaments can be explained by the distinct molecular mechanisms of action of inhibitor treatments. MLi-2 treatment is known to stabilize LRRK2 in its closed-kinase conformation which in turn promotes filament formation around microtubules. These observations are in stark contrast to LRRK2 localization in cells treated with GZD-824 which stabilizes LRRK2’s kinase in its inactive conformation weakening association with microtubules^19,21^. These findings confirm that inhibitor-induced transitions of LRRK2^IT^ from its active to inactive kinase conformation modulates its interaction with microtubules, leading to a shift from extensive to minimal microtubule decoration in cells.

The physiological relevance of LRRK2 filamentation around microtubules remains controversial. Wild-type LRRK2 does not exhibit enriched microtubule localization, whereas several Parkinson’s disease–associated mutants (e.g., I2020T, R1441H/G/S/T) show robust microtubule association. Microtubule decoration by LRRK2^IT^ has not been studied in cell types that endogenously express high levels of LRRK2, such as lung epithelial cells and brain-resident immune cells including microglia and macrophages^44^. Thus, it remains possible that aberrant LRRK2–microtubule interactions occur under physiological expression conditions, potentially disrupting homeostatic intracellular transport and being further exacerbated by type I LRRK2 inhibitors, as suggested by in vitro studies^23,45^.

In our previous work, we used in-cell cryo-ET to build and integrative model of the microtubule-associated catalytic ROC, COR, kinase and WD40 (RCKW) domains of LRRK2 in its closed-kinase conformation^22^. However, our model could not resolve the conformation of N-terminal LRR, ANK and ARM domains. Full-length LRRK2 has been captured in vitro in its intermediate active conformation^19–21,23–26^. However, how the N-terminal repeats of LRRK2 are organized when the protein is in its closed-kinase conformation remained unresolved. Stabilization of LRRK2 in a closed-kinase conformation by MLi-2 treatment and microtubule association reduces conformational heterogeneity to permit structure determination of full-length LRRK2^IT^ with the N-terminal repeats undocked from the catalytic core. Therefore, the key novelty of this structure is that it captures full-length LRRK2^IT^ in a cellular, microtubule-associated closed-kinase state and shows that kinase closure is compatible with an undocked N-terminal architecture. This distinguishes the in situ closed-kinase state from previously described in vitro intermediate active states. This structure represents one stabilized closed-kinase state within a broader conformational ensemble of undocked N-terminal repeats.

Our results clarify the relationship between kinase conformation, repeat undocking, and microtubule association. Increased microtubule association observed for I2020T mutant favors undocking of the N-terminal repeats, a prerequisite for kinase closure and filament assembly. In contrast, the G2019S mutation enhances catalytic activity without increasing microtubule association. Microtubule binding therefore reflects conformational state rather than kinase activity per se.

Together, these findings provide a structural view of full-length LRRK2 in a closed-kinase conformation and capture a resolved snapshot along its conformational continuum. Our results underscore how pathogenic mutations and distinct inhibitor classes reshape its structural landscape. More broadly, this work demonstrates the power of in-cell structural biology to illuminate dynamic regulatory transitions that are otherwise obscured by conformational heterogeneity.

## Author contributions

Conceptualization: T.B., E.V.

Sample Preparation and Data Acquisition: T.B., J.H., S.C., E.P.K., A.F.

Analysis: T.B., J.H.

Writing: T.B., A.E.L., E.V. with input from all the authors.

## Data Availability

Cryo-EM density maps have been deposited in the Electron Microscopy Data Bank under accession numbers EMD-77327 for Map A, corresponding to the focused map of the COR:COR interaction interface in LRRK2^I2020T^ + MLi-2 + microtubules; EMD-77328 for Map B, corresponding to the focused map of the WD40:WD40 interaction interface in LRRK2^I2020T^ + MLi-2 +microtubules; EMD-77329 for the large-lattice map of LRRK2^I2020T^ + MLi-2 + microtubules; and EMD-77330 for the LRRK2^I2020T^ + GZD-824 + microtubule map. The atomic model rigid-body docked into our maps is available under PDB accession code 8TZH.

## Acknowledgements

Electron microscopy data were collected at the UCSD Cryo-Electron Microscopy Facility, part of the Goeddel Family Technology Sandbox. We thank the UCSD Physics Computing Facility for computational support. This research was funded by Aligning Science Across Parkinson’s (grant number ASAP-000519) (A.E.L. and E.V.) through the Michael J. Fox Foundation for Parkinson’s Research (MJFF) (A.E.L. and E.V.), as well as from the National Science Foundation grant DBI 1920374 (to E.V.). T.B. was supported by an American Heart Association Predoctoral Fellowship. J.H. was supported by an EMBO long-term postdoctoral Fellowship (ALTF 871-2020). S.C. was supported by Jane Coffin Childs Memorial Fund for Medical Research. E.P.K. was supported by Jane Coffin Childs Memorial Fund for Medical Research. E.V. is a Howard Hughes Medical Institute Investigator.

## Methods

### Cell Culture

293T cells were cultured in Dulbecco’s Modified Eagle’s Medium (DMEM) formulated with high glucose. DMEM was supplemented by 10% (v/v) heat inactivated fetal bovine serum (FBS), 0.1mM Non-Essential Amino Acids (NEAA), 1mM Sodium Pyruvate, 1% penicillin-streptomycin (Pen-Strep), 500µg/ml Geneticin Selective antibiotic. Cells were maintained at 37°C and 5% CO_2_ subcultured every 2-3 days at ∼80% confluency. Cells were detached for subculture using 0.05% trypsin-EDTA.

### Transfection Experiments

293T cells were transfecting using the Lipofectamine 3000 Transfection Kit (Thermo Fisher Scientific) according to the manufacturer’s protocol. For each transfection, 1µg of GFP-tagged LRRK2 recombinant DNA was used (Addgene). Transfections were carried out in 6-well plates. Cells were incubated for 48 hours post-transfection at 37°C and 5% CO_2_. Prior to sample preparation for fixation or cryo-EM grid preparation cells were treated with 500nM of MLi-2 for 2 hours, or 300nM GZD-824 for 10-30 minutes. Following treatment, transfected cells were either fixed for fluorescence microscopy analysis or processed for cryo-ET grid preparation.

### Cryo-TEM Grid Preparation

Au Quantifoil R1/4 200 mesh grids were used to prepare cellular samples. Prior to cell deposition the grids were patterned to enhance cell adhesion and positioning for cryo-ET^46^. In brief, grids were plasma cleaned and were treated with PLL-PEG. Using UV laser custom defined micropatterns were created in the layer of PLL-PEG creating adhesive regions using the PRIMO micropatterning system (Alveole). The patterned grids were incubated with fibronectin for 1 hour at room temperature before cells were deposited on them. Detailed protocol for patterning grids can be accessed here^47^.

Cells were detached from culture dishes after transfection via trypsin treatment, counted and approximately 200,000 cells were deposited on four grids in a 3cm cell culture dish. The cells were then incubated on the patterned grids for 2 hours or until the cells were completely attached.

Following attachment to the grids, samples were plunge frozen using a manual custom plunger assembly. The freezing medium consisted of 50/50 ethane/propane mixture. Grids were blotted for 6-8 seconds in a humidity-controlled room. Plunge frozen grids were secured in autogrids and stored in liquid nitrogen until imaging.

### Cryo-Focused Ion Beam Milling

Cryo-FIB milling was performed on an Aquilos 2+ system from Thermo Fisher Scientific. Scanning electron microscope images were taken using either 2KV or 5KV at a 30pA beam current. The lamella were prepared at 11° stage angle. Milling was performed in sequential manner starting from rough milling with 0.5nA-0.1nA currents, followed by intermediate milling at 50-30pA and final polishing using 10pA current. Target lamella thickness was 150-200nm. A detailed protocol for cryo-FIB milling is described in previous publications^41,48,49^.

### Integrated Cryo-Fluorescence Microscopy

Cryo-Fluorescence microscopy was performed using an integrated fluorescence microscopy (iFLM) system on Aquilos 2+. The 20x air objective NA 0.7 was used for imaging. For GFP tagged cellular samples excitation 470nm was used for visualization. Illumination was set to 10% intensity with 200ms exposure time to minimize photodamage. Z-stacks spanning 10-20µm in depth were acquired with step sizes ranging between 500-800nm. This allowed imaging of the whole cellular volumes with adequate resolution and limited photodamage.

### Cryo-Electron Tomography

TEM image acquisition was performed on Titan Krios G3 system equipped with a K3 direct electron detector and Gatan energy filter. The milling slot of the autogrid was oriented perpendicular to the loading cartridge. Tilt-Series were acquired using Serial-EM software and Pacetomo automated tilt-series acquisition methods^50,51^. The images were collected at a target defocus range of 4-5 µm with a pixel size of 1.341 Å. Dose symmetric data acquisition schemes were implemented with a tilt range of ±54° degrees with 3° increments^52^. The pre-tilt was calculated using Pacetomo and acquisition was initiated from this position. Cumulative electron dose of approximately 140 e^−^/Å^2^ was applied across the tilt series.

### Tomogram Reconstruction and Annotation

Tilt-series were processed using a Warp-Imod pipeline^53–55^. Initial preprocessing including frame alignment, gain correction, and motion correction was performed in Warp. Tilt-series were aligned in IMOD. The aligned tilt-series stacks were imported back into Warp 1.0.9/beta for 3D-CTF correction. Final tomograms were reconstructed in Warp and visualized in IMOD.

### Tomogram Segmentation and Visualization

Cellular membrane segmentation was performed using MemBrainSeg v0.0.9 with v10_alpha pretrained segmentation model with manual clean-up performed in UCSF ChimeraX^56,57^. Microtubule coordinates were traced manually in IMOD and visualized in ChimeraX. Ribosomes were identified through template matching in Warp using an ab initio ribosome model, with subsequent coordinate refinement in Relion. LRRK2 subtomograms were extracted from Dynamo coordinate tables^58^. The segmented membranes, microtubules, LRRK2 and ribosome coordinates were inputted in ChimeraX and ArtiaX for comprehensive visualization^59^.

### Microtubule tracing and protofilament number analysis

Tomograms were visualized in IMOD to identify LRRK2^IT^ decorated microtubules. These microtubules were manually traced in IMOD and coordinates were saved as contours. Filaments were cropped equidistantly along the traced microtubule coordinates using filament path function in Dynamo. The cropped volumes were averaged and smoothed to generate an ab initio reference for a microtubule. Based on this ab initio reference a cylindrical featureless reference was generated. Each microtubule was cropped, and alignment was run to symmetrize each individual microtubule. The results of the alignment were evaluated to assess the protofilament number of the microtubule.

### LRRK2 subtomogram analysis

LRRK2 decorated microtubule coordinates were used to extract subtomograms of LRRK2 using Dynamo filament path function. 15 subunits at 8A separation were used to crop the LRRK2 subtomograms along the microtubule path. Cropped particles were averaged to generate initial reference. Spherical mask around central 4 subunits of LRRK2 was generated and initial refinements were run in Dynamo to align cropped particles. We removed duplicate particles from these subtomograms and exported them for import into Relion3. The subtomograms were refined in Relion3 with a C1 symmetry^60^. Focused refinements were performed on four central subunits of LRRK2^IT^. 3D classification was performed following initial refinements and classes with distinct LRRK2 decorations were selected for further refinements. The refinements were either centered on COR-COR interface of the protein or WD40-WD40 interaction interface. The center of the box was shifted accordingly, and refinements were performed with newly extracted particles.

### Model Building

Structures of previously published LRRK2 were used as references for model building. ChimeraX rigid body fitting was used to fit each individual domain into the density maps. The fitting was performed stepwise. The domains of WD40and kinase C-lobe were separated and fit into the maps together. Following this, kinase N-lobe, COR and ROC domains were fitted into the density for the LRRK2. LRR-ANK domains were fit separately as well accounting for the remaining electron density in the in-situ maps. ARM domain was obtained through Alphafold2 model of LRRK2 and separated from the rest of the protein. ARM domain was individually mapped into the density for the N-terminal domains in electron density maps with larger lattice of LRRK2 accounting for the extra densities.

