## Supplemental_material for "In-cell cryo-electron tomography reveals differential effects of type I and type II kinase inhibitors on LRRK2 filament formation and microtubule association"

bioRχiv

### Extended Data Figures 1-8

### Tables 1-2

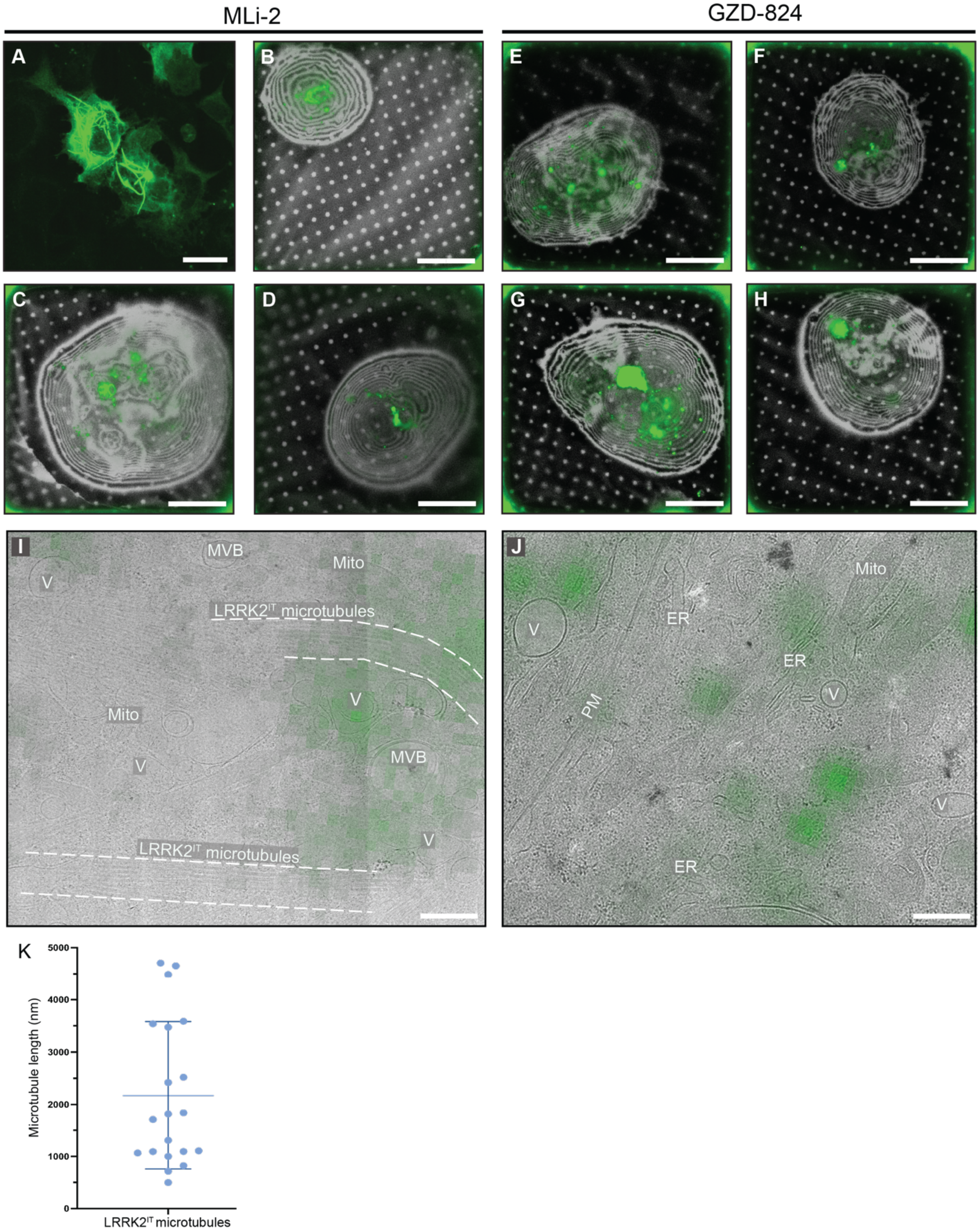

**Extended data figure 1. LRRK2$^{IT}$ expression in 293T cells responds differently to treatment with type I and type II kinase inhibitors.** **(A)** LRRK2$^{IT}$ microtubule-associated filaments (green) in 293T fixed cells treated with MLi-2. **(B-D)** Cryo-fluorescence micrographs of GFP-LRRK2$^{IT}$ -expressing cells treated with MLi-2 show a distribution of LRRK2$^{IT}$ as filaments, puncta and diffuse in the cytosol. **(E-H)** Cryo-fluorescence microscope images of GFP-LRRK2$^{IT}$ expressing cells treated with GZD-824 displaying diffuse and punctate distribution of LRRK2$^{IT}$. **(I-J)** Cryo-TEM lamella overview LRRK2$^{IT}$ expressing cell treated with either MLi-2 (I) or GZD-824 (J). GFP-LRRK2$^{IT}$signal (green) is overlayed from cryo-FM imaging to guide localization of LRRK2$^{IT}$. **(K)** Length of LRRK2$^{IT}$ filaments evaluated from the cryo-TEM overview in (I) in cells treated with MLi-2. Each data point represents a single filament. (22 data points; vertical bar represents mean with standard deviation 2.17±1.4μm). MVB: Multivesicular body V: Vesicles, ER: Endoplasmic reticulum, Mito: Mitochondria, PM: Plasma membrane. Scale bars: A 40μm; B-H 20μm; I-J 500nm.

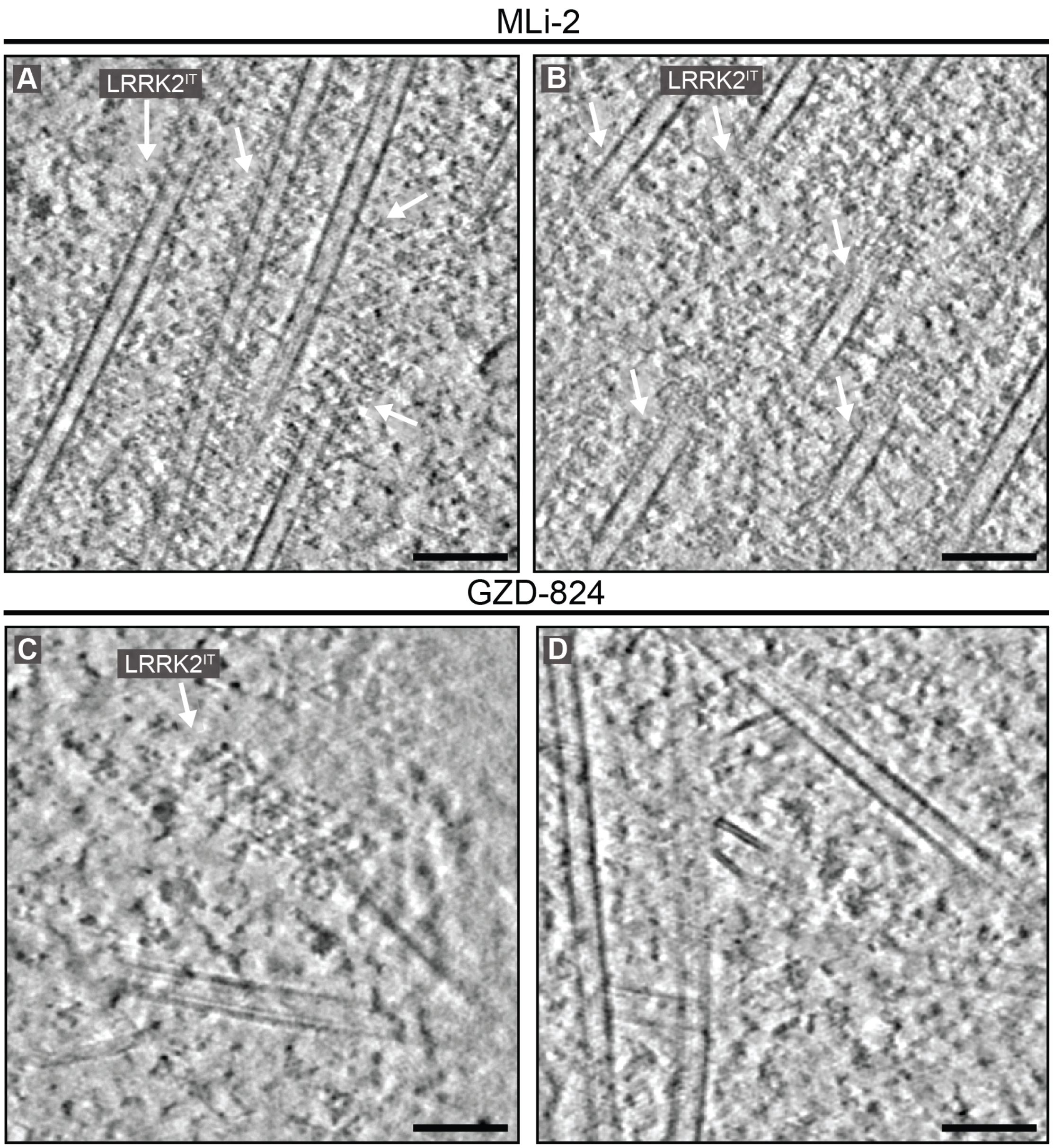

**Extended data figure 2. LRRK2$^{IT}$ filament formation around microtubules is extensive when treated with type I, but not type II inhibitor. (A-B)** Representative tomogram slice through the cytosol highlighting multiple LRRK2$^{IT}$ decorated microtubules in a cell treated with MLi-2. **(C-D)** Tomogram snapshots of a cell treated with GZD-824 where majority of microtubules are not decorated with LRRK2$^{IT}$. Scale bars: A-D 100nm.

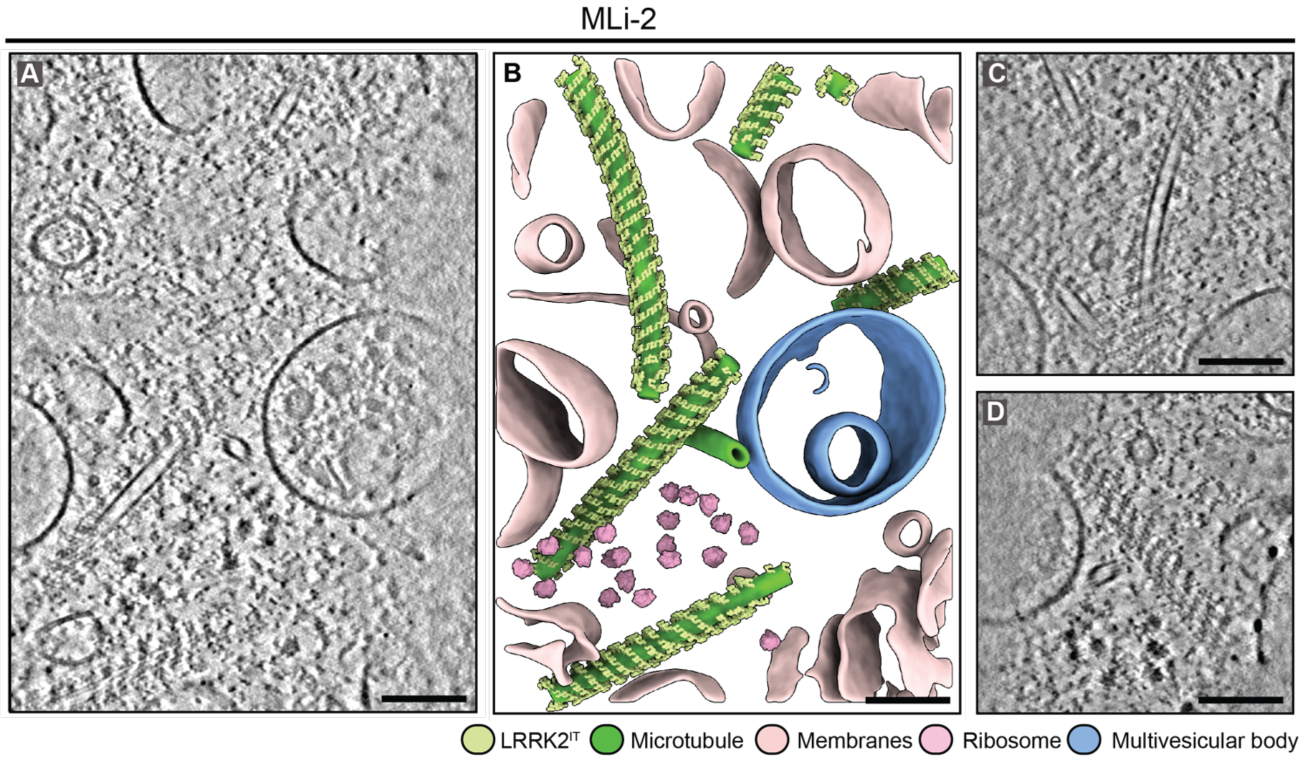

**Extended data figure 3. Type I inhibitor treated cells have single microtubules decorated with LRRK2$^{IT}$**

**(A)** Tomogram slice of a cell treated with MLi-2 showing six single non-bundled microtubules decorated with LRRK2$^{IT}$. **(B)** Segmentation of the tomogram shown in A highlighting LRRK2$^{IT}$ filamentation around microtubules. **(C, D)** Tomogram snapshot of a LRRK2$^{IT}$ lattice in MLi-2 treated cell. Lime Green: LRRK2$^{IT}$, green: microtubule, light pink: vesicles, blue: multivesicular bodies, pink: ribosomes. Scale bars: A-D 100nm.

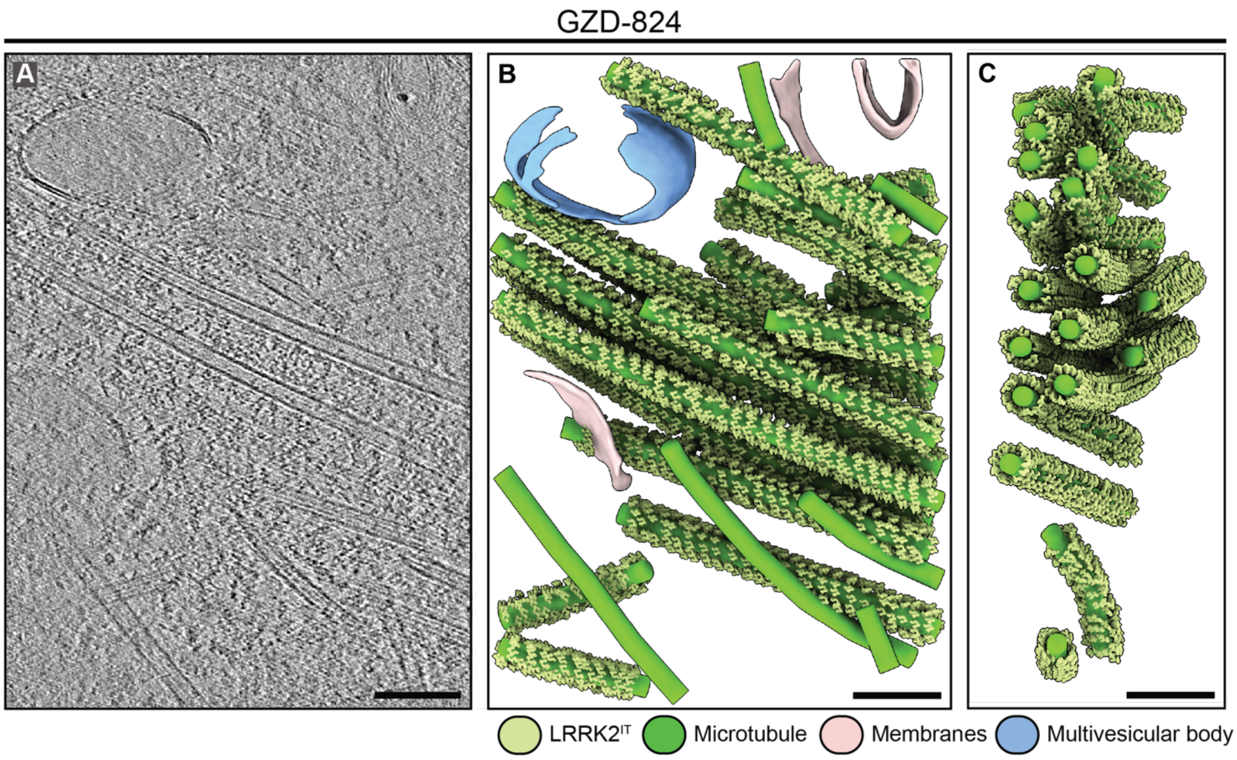

**Extended data figure 4 A single instance of LRRK2$^{IT}$ decorated microtubules was captured in cells treated with Type II inhibitor.**
**(A)** Tomogram snapshot of a cell treated with GZD-824 showing a LRRK2$^{IT}$ decorated microtubule bundle. This is the single instance LRRK2$^{IT}$ bundle captured in cells treated with GZD-824. **(B)** Segmented representation of the tomogram shown in A. **(C)** Side view of the LRRK2$^{IT}$ decorated microtubule bundle shown in B. Lime Green: LRRK2$^{IT}$, green: microtubule, light pink: vesicles, blue: multivesicular bodies. Scale bars: A-C 100nm.

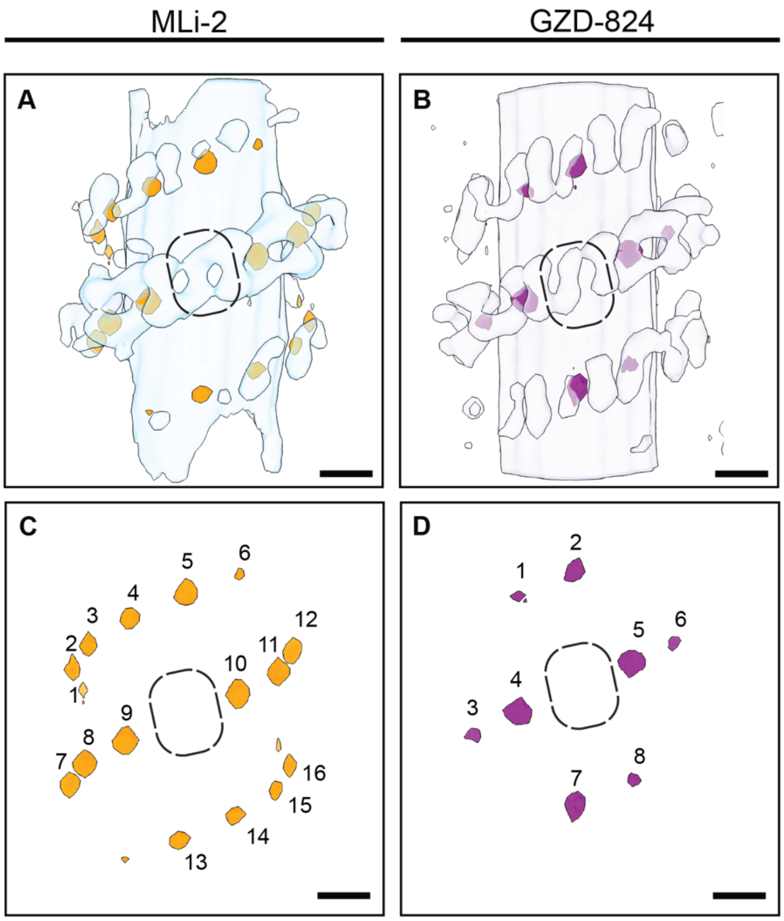

**Extended data figure 5. Nearest neighbor analysis of a central LRRK2[IT] subunit**

**(A-B)** Nearest neighbor analysis of a central LRRK2[IT] subunit (highlighted in black) in MLi-2 and GZD-824 treated cells. **(C-D)** Annotation of immediate neighboring LRRK2[IT] subunits within each lattice. In MLi-2 treated cells, the central LRRK2[IT] subunit is surrounded by 16 nearest neighbors highlighted in orange. While in GZD-824 treated cells, the central LRRK2[IT] subunit has only 8 nearest neighbors shown in pink.

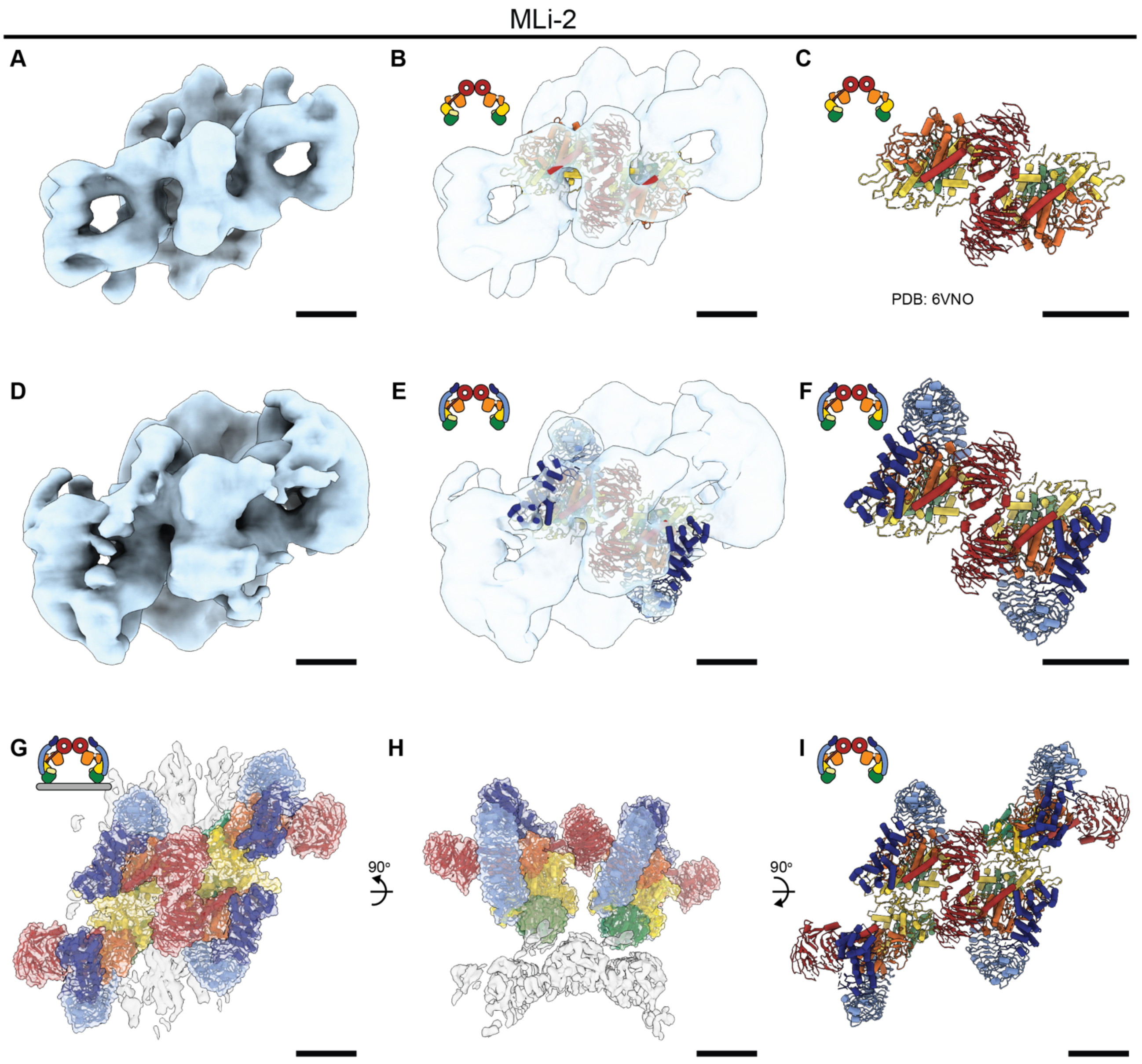

**Extended data figure 6. Subtomogram analysis of LRRK2$^{IT}$ in MLi-2 treated cells centered on WD40-WD40 domain interaction interface.**

**(A)** Subtomogram average map of LRRK2$^{IT}$ centered on the WD40-WD40 domain interface. (threshold value: 0.12) **(B, C)** Molecular model of the closed-kinase LRRK2 fit in the map in A demonstrating interaction interface between WD40-WD40 domains. Domain architecture color scheme is the same as given in Fig. 1A. **(D)** Subtomogram average map at threshold value 0.24 showing the densities for N-terminal domains of LRRK2$^{IT}$. **(E, F)** Molecular model of full-length LRRK2 fit in the map in D showing the architecture of the N-terminal domains. **(G-I)** Molecular model of the four LRRK2 protomers fit into subtomogram map shown in D demonstrating overall architecture of the full-length LRRK2$^{IT}$ assembly around a microtubule. Scale bars: A-I 5nm.

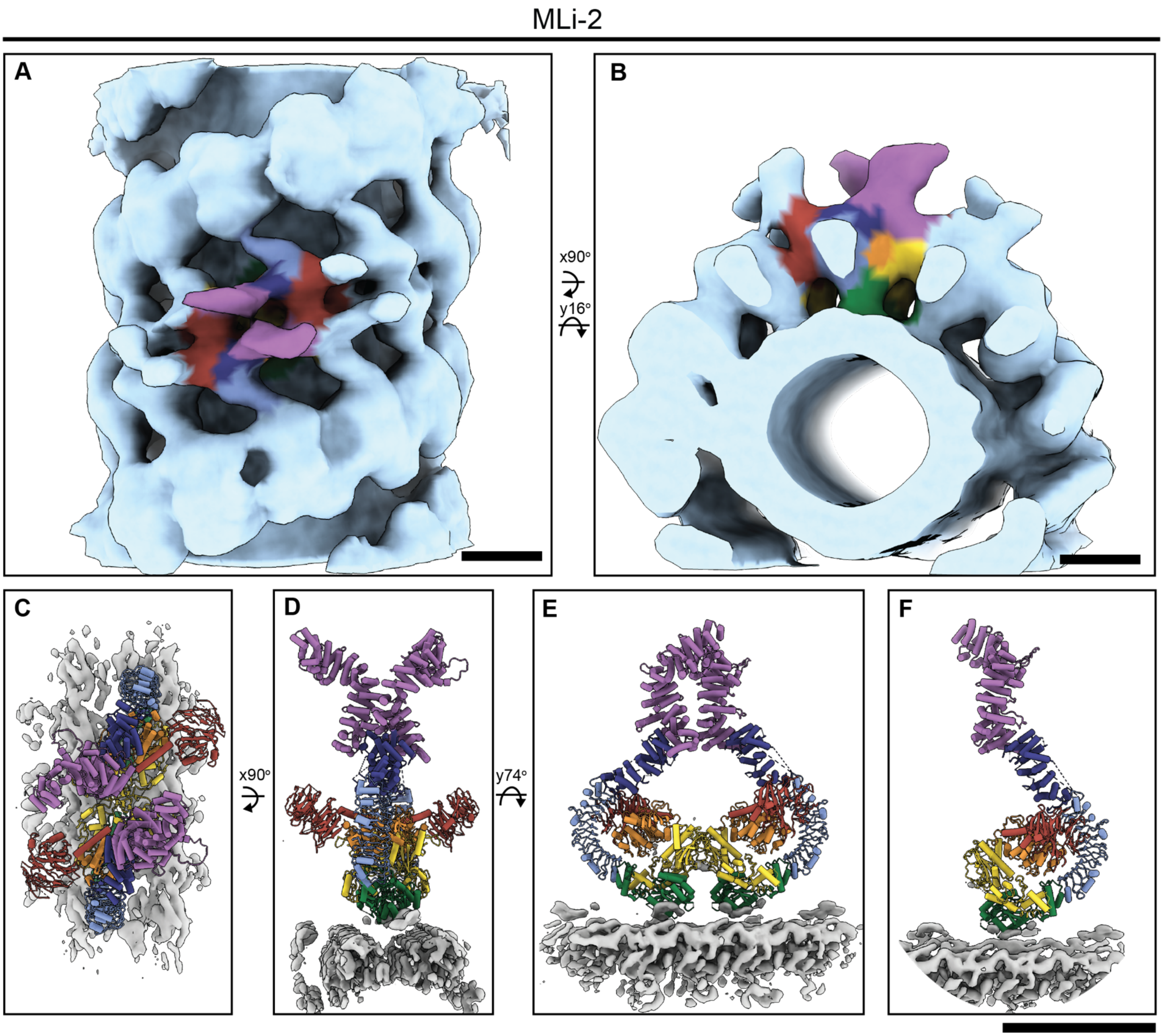

**Extended data figure 7. Model of full-length LRRK2$^{IT}$ associated with microtubules in its active-like conformation.**
**(A-B)** Subtomogram average of LRRK2$^{IT}$ lattice decorating a microtubule and a corresponding cross-section. **(C-E)** Molecular model of the closed-kinase LRRK2 fit in the map in A demonstrating domain organization of the N-terminal LRR-ANK-ARM domains. **(F)** Molecular model of a single full-length LRRK2 fitted into the map shown in A, highlighting the architecture of the N-terminal domains. Domains are colored in as in Fig 1A, microtubule is presented in gray (EMDB 25908). Scale bars: A-F 10nm.

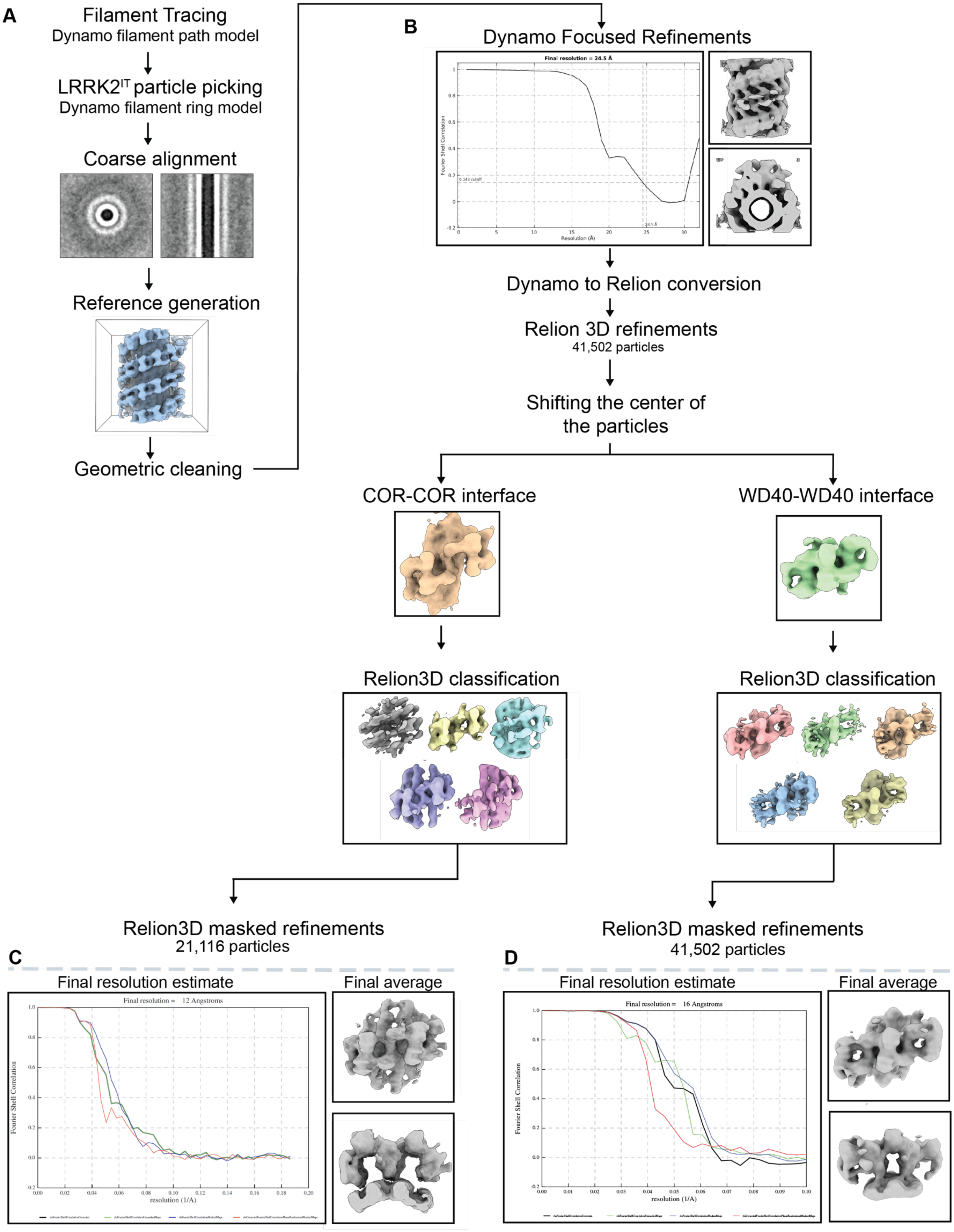

A
Filament Tracing
Dynamo filament path model
LRRK2^IT particle picking
Dynamo filament ring model
Coarse alignment
Reference generation
Geometric cleaning
B
Dynamo Focused Refinements
Final resolution = 24.5 Å
Fourier Shell Correlation
0.143 cutoff
24.5 Å
Resolution (Å)
Dynamo to Relion conversion
Relion 3D refinements
41,502 particles
Shifting the center of
the particles
COR-COR interface
WD40-WD40 interface
Relion3D classification
Relion3D classification
Relion3D masked refinements
21,116 particles
Relion3D masked refinements
41,502 particles
C
Final resolution estimate
Final average
Final resolution = 12 Angstroms
Fourier Shell Correlation
resolution (1/A)
D
Final resolution estimate
Final average
Final resolution = 16 Angstroms
Fourier Shell Correlation
resolution (1/A)

**Extended data figure 8. Data-processing workflow for LRRK2$^{IT}$ in type I inhibitor treated cells.**

**(A)** Cryo-ET data processing workflow for LRRK2$^{IT}$ decorated microtubules in MLi-2 treated cells. **(B)** Fourier shell correlation (FSC) curve for the Dynamo-refined MLi-2–treated LRRK2 subtomogram average presented in Figure 5. **(C)** Final maps and the gold standard Fourier Shell Correlation (FSC)curves (0.143 cutoff) show the final resolution of the COR-COR domain interaction interface centered LRRK2$^{IT}$ refinement **(D)** Final maps and the gold standard Fourier Shell Correlation (FSC)curves (0.143 cutoff) show the final resolution of the WD40-WD40 domain interaction interface centered LRRK2$^{IT}$ refinement.

**Table 1. List of key reagents and tools used to perform this study.**

| Reagent or Resource | Source | Catalog Number / Identifier |
|---|---|---|
| Chemicals, Peptides, and Recombinant proteins | | |
| MLi-2 | Tocris | 5756 |
| Taxol (paclitaxel) | Cell Signaling Technology | 9807 |
| GZD-824 | Cayman Chemical | 21508 |
| DNL201 (GNE0877) | MedChem Express | HY-15796 |
| Poly-L-Lysine solution 0.1% (w/v) in $H_2O$ (PLL-PEG) | Millipore sigma | P8920 |
| Fibronectin | Thermo Fisher Scientific (Gibco) | 33016015 |
| Fetal Bovine Serum (FBS) | Thermo Fisher Scientific (Gibco) | A5209402 |
| MEM NON-Essential Amino Acis (NEAA) | Thermo Fisher Scientific (Gibco) | 11140050 |
| Geneticin | Thermo Fisher Scientific (Gibco) | 10131035 |
| Pen-Strep | Thermo Fisher Scientific (Gibco) | 15140122 |
| Sodium Pyruvate | Thermo Fisher Scientific (Gibco) | 11360070 |
| DMEM (high-glucose) | Thermo Fisher Scientific (Gibco) | 11965092 |
| DMSO, Anhydrous | Thermo Fisher Scientific | D12345 |
| Trypsin-EDTA 0.25% | Thermo Fisher Scientific | 25200056 |
| Critical Commercial Assays | | |
| Lipofectamine 3000 Transfection Reagent | Thermo Fisher Scientific | L3000001 |
| Deposited Data | | |
| Cell Line | | |
| HEK293FT cells | Thermo Fisher Scientific | R70007 |
| Recombinant DNA | | |
| GFP-LRRK2-WT | Addgene | 25044 |
| GFP-LRRK2-I2020T | Watanabe et al. 2021 | |
| GFP-LRRK2-G2019S | Addgene | 25045 |
| Software and Algorithms | | |
| TFS TUI for Aquilos | Thermo Fisher Scientific | - |
| MAPS 3.24 | Thermo Fisher Scientific | - |
| iFLM V2 | Thermo Fisher Scientific | - |
| Serial EM | Mastronarde, 2005 | - |
| Pace-Tomo | Eisenstein et al., 2023 | - |
| Warp 1.0.9 / beta 1 | Tegunov et al., 2021, Tegunov et al., 2019 | - |

| | | |
|---|---|---|
| IMOD | Kremer et al., 1996 | - |
| MemBrainSeg | Lamm et al., 2024 | - |
| ChimeraX | Meng et al., 2023 | - |
| ArtiaX | Ermel et al., 2022 | - |
| DYNAMO | Castano-Diez et al., 2012 | - |
| RELION | Bharat et al., 2015 | - |
| Other | | |
| Au Quantifoil R1/4 grids 200 mesh | Quantifoil Micro tools | NA |
| #1 Whatman filter paper | Whatman | 1001 |
| Liquid Nitrogen | Airgas | - |
| Ethane-Propane mixture | Airgas | - |

**Table 2. Data acquisition table for cryo-ET datasets used in this study.**

| Parameters | Data # MLi-2 | Data # GZD-824 |
|---|---|---|
| Magnification | 63,000 | 63,000 |
| Voltage (kV) | 300 kV | 300 kV |
| Total dose ($e^{-}/Å^{2}$) | ~140 $e^{-}/Å^{2}$ | ~140 $e^{-}/Å^{2}$ |
| Defocus range (μm) | -2 to -5 | -2 to -5 |
| Acquisition scheme | Dose symmetric | Dose symmetric |
| Pixel size (Å) | 1.3410 Å | 1.3410 Å |
| No. of frames | 6 | 6 |
| # of tomograms | 26 | 6 |
| # of LRRK2$^{IT}$ decorated microtubules | 254 | 27 |
| Final particle number | 21,116 particles | 6,212 particles |
| Symmetry imposed | C2 | C1 |
| Map resolution (Å) | ~12 Å | ~30 Å |
